# Direct fast estimator for recombination frequency in the F_2_ cross

**DOI:** 10.1101/2021.04.20.440219

**Authors:** Mathias Lorieux

**Affiliations:** DIADE, University of Montpellier, Cirad, IRD, Montpellier, France

**Keywords:** Recombination fraction, F_2_ cross, EM algorithm, efficient estimator

## Abstract

In this short note, a new unbiased maximum-likelihood estimator is proposed for the recombination frequency in the F_2_ cross. The estimator is much faster to calculate than its EM algorithm equivalent, yet as efficient. Simulation studies are carried to illustrate the gain over another simple estimate proposed by Benito & Gallego (2004).

## 1 Introduction

Whole-genome sequencing allows constructing genetic maps with millions of loci. Unless the order of loci is known *a priori*, determining the order of the loci on linkage groups requires computing the recombination fractions for all pairs of loci, in the order of trillions. Computing time can thus be extremely long, especially in the F_2_ cross where an iterative procedure is usually used by most of the genetic mapping software. A faster yet efficient estimator is proposed, which can be used as replacement for the iterative methods.

## 2 Methods

Consider two loci for which we want to know their recombination frequency *r*. In the F_2_ cross, it is not possible to count the recombinant individuals directly, since the double heterozygotes class is issued from either two recombinant *(AB*|*BA* or *BA*|*AB*) or two non-recombinant (*AA*|*BB*) gametes, if *A* and *B* are the parental alleles. Those two configurations are generally indistinguishable (see Appendix 1 for a detailed explanation). Thus, we cannot estimate *r* directly since we need to solve the 3rd degree equation to obtain the efficient, unbiased maximum-likelihood estimator for *r*

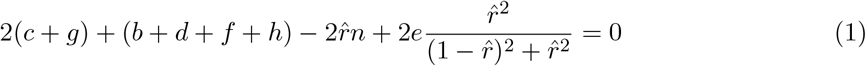

where *c* and *g* are the counts for the double recombinants *AB*|*AB* or *BA*|*BA, b, d, f* and *h* are the counts for the single recombinants *AA*|*AB, AB*|*AA, AB*|*BB* and *BB*|*AB*, and *e* is the count for the double heterozygotes class. Note that the non-recombinant classes *AA*|*AA* and *BB*|*BB, a* and *i*, are not used since they are not informative for the estimation of *r*.

The common way to solve (1) is to use an iterative algorithm to estimate the fraction of the recombinants, *AB* |*BA* or *BA*|*AB*, in the double heterozygote class. The most used in this case are the Newton-Raphson (see for instance Edwards 1972) and Expectation-maximization (EM) algorithms (Dempster et al, 1977). For instance, an EM algorithm to estimate the fraction of double recombinants in the *e* class was proposed by Mangin (1991)

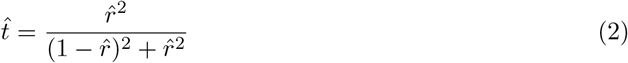

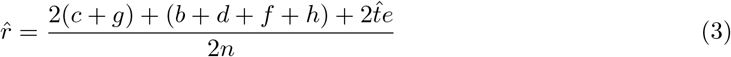

After convergence, we obtain the 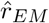 estimator, which asymptotic variance is

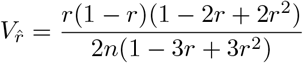

Iterative algorithms can require many iterations to reach convergence, depending on the starting parameter value for *r*, making them slow to calculate. Benito and Gallego (2014) proposed a simplified, faster estimator for *r* (here noted as 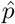) that ignores the double heterozygotes class. Following our notations, this estimator is

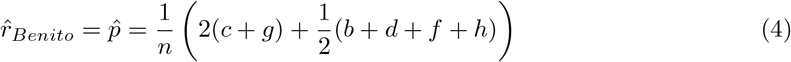

With asymptotic variance

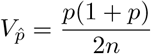

This estimator is unbiased. However, it has greater variance than the 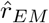 estimator, the ratio 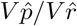 being

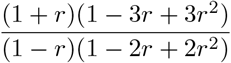

This ratio increases with *r*, which is problematic for testing linkage *(H*_*0*_ : *r* = 0.5).

Now, note that if a value close enough to the true *r* is chosen to initiate the EM process, then convergence is reached in very few steps. Even the first step gives a better estimation than the initialization value. We can thus try to reduce the EM process to a direct estimator for *r*, that is, without having to iterate. One can use the unbiased Benito’s estimator, 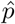, as the starting value in a one-step EM process. For this, we replace 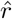 in the 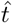 EM formula (2) by 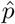

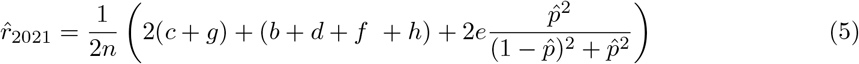

then we plug in (4) in (5)

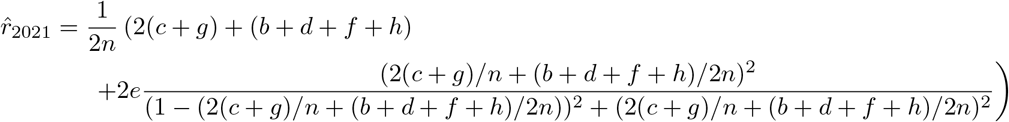

Let *R* = *c* + *g* and *H* = *b* + *d* + *f* + *h* in (4). Then

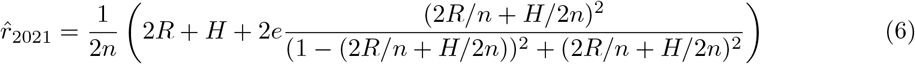

The fastest way to compute 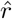 may depend on the programming language. However, multiplication is usually faster than division in computers, so one might want to pre-calculate *R, H, D* = 1/2*n, E* = 4*DR* + *DH* and *F* = *E*^2^ then apply the formula

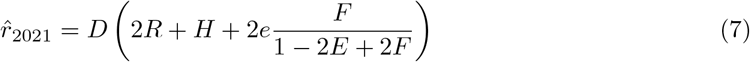

An estimator’s efficiency is obtained by calculating its mean-squared error (MSE), which is the sum of its variance and the squared difference between its estimated and expected values (i.e., its bias). The smaller the MSE, the greater the efficiency. Note that if an estimator is unbiased, the asymptotic MSE is simply the variance, however in a finite number of observations it is preferable to keep the observed bias in the MSE formula. Monte-Carlo simulation studies were carried to verify the properties of 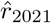 compared with 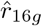 (defined in Appendix 1), 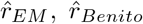. For this, for each value of *r* (0-0.5, step 0.01) 1,000,000 F_2_ populations were simulated and the four estimators calculated. Convergence of 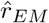 was stopped using 0.00001 as stop criterion to ensure comparable precision with the two other estimators, and 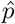 as initialization parameter.

Computing times for 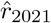 and 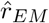 were compared by the same simulation strategy, this time varying the initialization parameter (0-0.5, step 0.01). The only difference is that 500,000 populations were used.

In Supplemental data 1, an Excel workbook is provided to replicate the simulation studies with any population size, number of repetitions and value for *f*.

## 3 Results

The new estimator 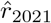 is equivalent to 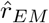 after one iteration as it is unbiased and has approximately the same asymptotic variance.

Figure 1 shows plots of the MSEs and standard deviations of four estimators for 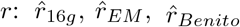 or 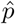 and formula (7) 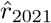 in simulated F_2_ populations. We see that the relative efficiency of the new estimator over Benito and Gallego’s 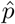 increases continuously with *r*.

**Figure 1:**
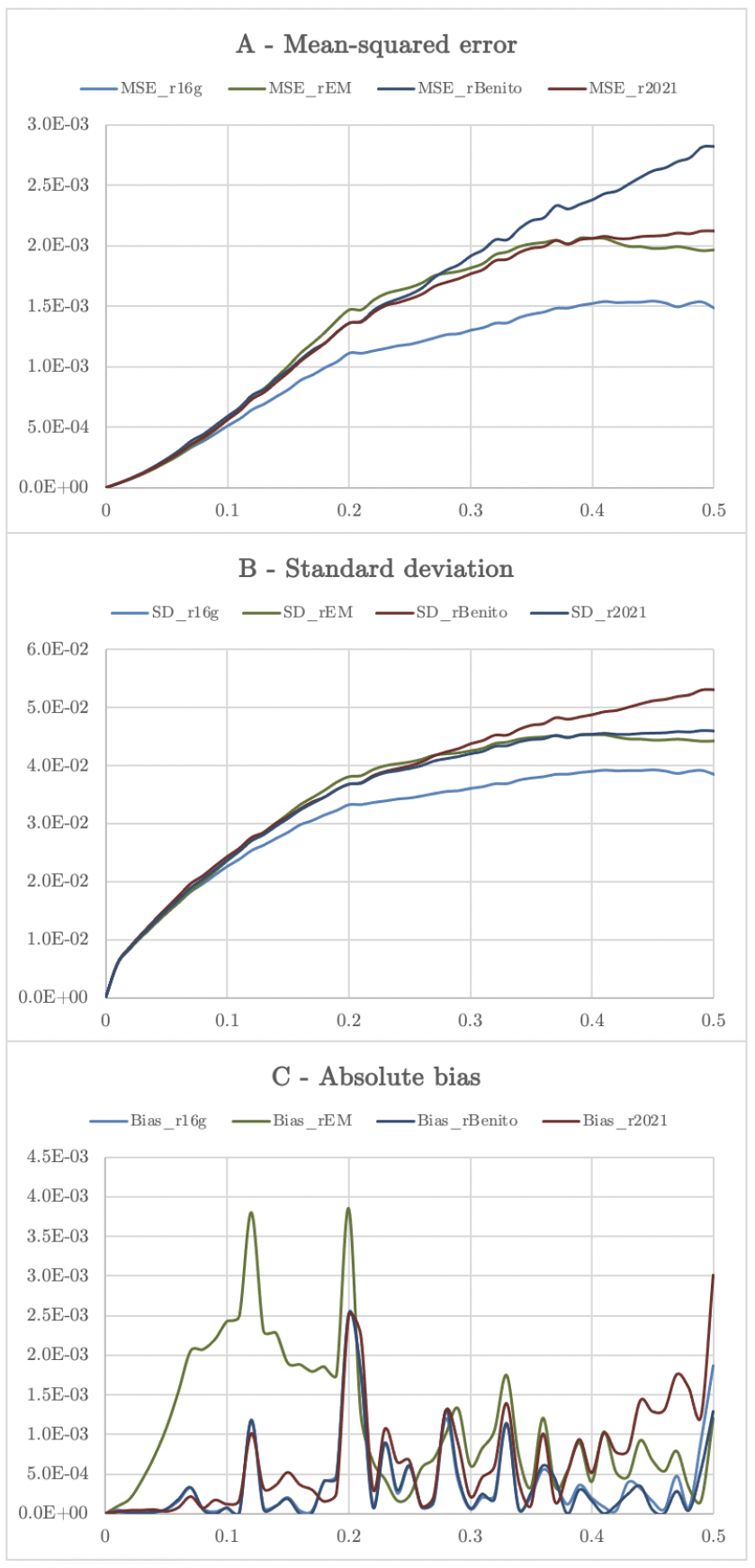
Mean-squared errors and standard deviations of three estimators for *r*, obtained by Monte-Carlo simulation of 1,000,000 F2 populations of size *n =* 100. Horizontal axis: expected recombination frequency *r*. A: Mean-squared errors for maximum-likelihood 16 genotypes (MSE_r16g, defined in Appendix 1), maximum-likelihood/EM (MSE_rEM) *r*, Benito and Gallego’s 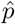 (MSE_rBenito) and Formula (7) (MSE_r2021). B: standard deviations. C: Absolute bias. Convergence of 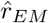 was stopped at 0.00001 and 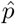 was used as initialization parameter.

Moreover, 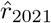 can be more precise than 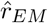 (Figure 1C), especially when an inadequate starting value is chosen and few iterations are allowed to speed-up the computation.

Regarding computing time, in the simulation study Formula (7) was 3 to 17.5 times faster than the EM algorithm estimator to compute in 95% of the cases (Figure 2). The relative computation time depended on both the values of *r* and the initialization value *r*_*Start*_. Computing time of course also depends on the stop criterion set for the convergence.

**Figure 2:**
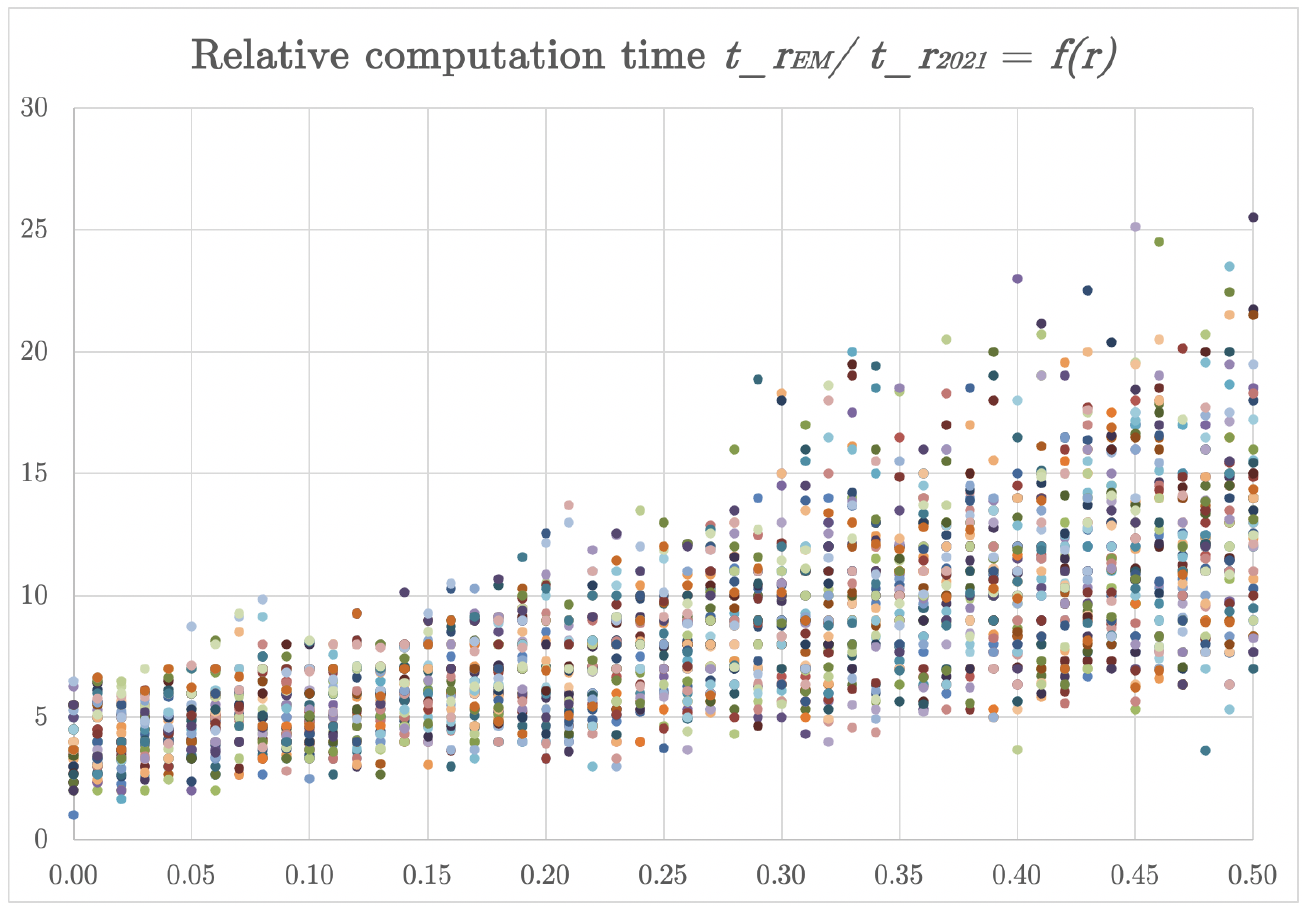
Relative computation time *t*_*r*_*EM =t*_*r*_*2021*, obtained by simulation of 500,000 F_**2**_ populations of size *n* = 100. Horizontal axis: expected recombination frequency *r*. Convergence of 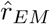 was stopped at 0.00001. Each dot color represents a different value for the initialization parameter, which varied between 0 and 0.5.

## 4 Discussion

The new proposed estimator, 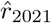, has the desirable properties of efficiency — it is unbiased and has minimal variance — and is considerably faster to compute than the EM algorithm. As 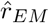, it is more efficient than Benito and Gallego’s estimator, especially for *r* values greater than 0.3.

Also, it should be noted that in case of segregation distortion that, for instance, would favor the double heterozygotes class *e*, Benito and Gallego’s estimator would rely on a smaller number of observations than expected and would have a larger standard deviation. This is not the case for 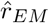 and 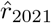.

Using the new estimator 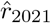 in lieu of *r*_*EM*_ in F_2_ crosses when the number of loci is very large could help researcher computing genetic maps in less time, especially in ”orphan” species that do not benefit from a reference sequence. 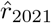 can also replace *r*_*Benito*_ in most cases.

## 6 Figures

## 7 Appendix 1

### Details of the estimation of recombination fraction in the F_2_ cross (two codominant loci, two alleles)

We consider the case of two loci segregating in an F_**2**_ progeny. If A and B are the parental alleles, 16 genotypes are produced in the F_**2**_ (Table A1).

**Table A1:** The true genotypes (female/male): 16 genotypes.

|  | Male<br>gamete | non-recombinant | non-recombinant | recombinant | recombinant |
| --- | --- | --- | --- | --- | --- |
| Female<br>gamete |  | AA | BB | AB | BA |
| non-recombinant | AA | AA\|AA | AA\|BB | AA\|AB | AA\|BA |
| non-recombinant | BB | BB\|AA | BB\|BB | BB\|AB | BB\|BA |
| recombinant | AB | AB\|AA | AB\|BB | AB\|AB | AB\|BA |
| recombinant | BA | BA\|AA | BA\|BB | BA\|AB | BA\|BA |

In pure lines crosses, there are only two possible alleles, so only 9 genotypes can be observed (Table A2).

**Table A2:** What we see: 9 genotypes.

|  | Male<br>gamete | non-recombinant | non-recombinant | recombinant | recombinant |
| --- | --- | --- | --- | --- | --- |
| Female<br>gamete |  | AA | BB | AB | BA |
| non-recombinant | AA | A\|A (1) | H\|H (5) | A\|H (2) | H\|A (4) |
| non-recombinant | BB | H\|H (5) | B\|B (9) | H\|B (6) | B\|H (8) |
| recombinant | AB | A\|H (2) | H\|B (6) | A\|B (3) | H\|H (5) |
| recombinant | BA | H\|A (4) | B\|H (8) | H\|H (5) | B\|A (7) |

In the double heterozygotes class *H*|*H*, the double recombinants cannot be distinguished from the non-recombinants (Table A3).

**Table A3:**
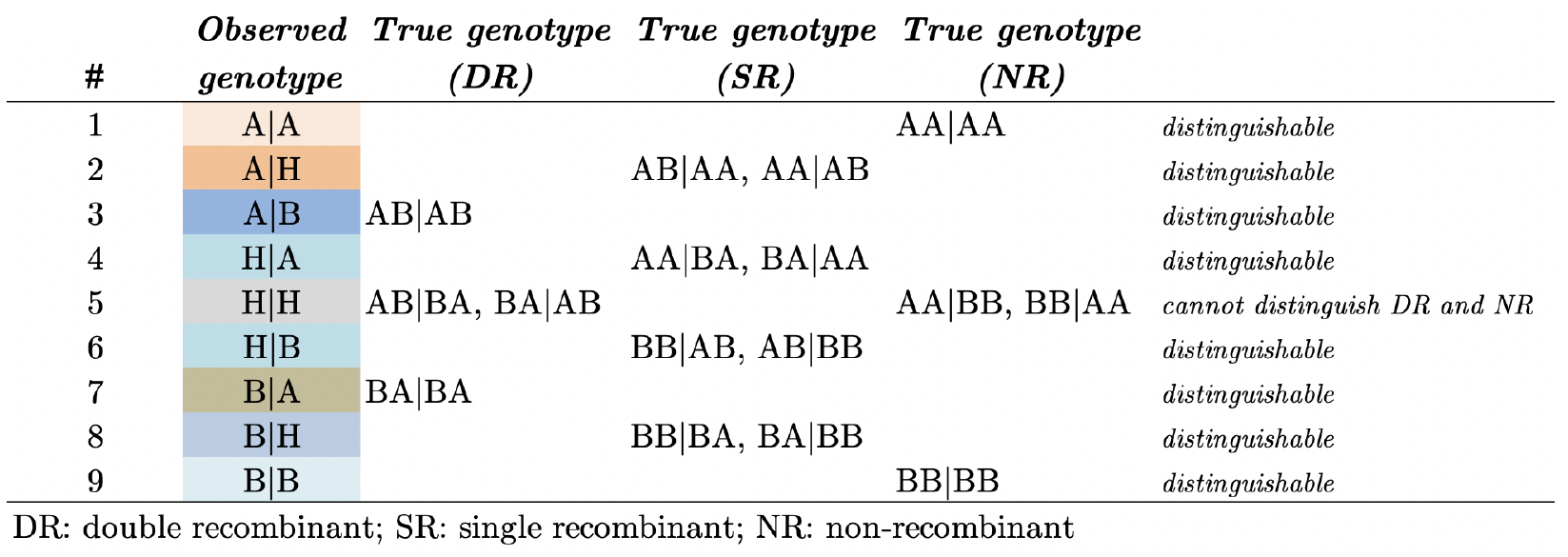
Observable vs. true genotypes.

| # | Observed<br>genotype | True genotype<br>(DR) | True genotype<br>(SR) | True genotype<br>(NR) |  |
| --- | --- | --- | --- | --- | --- |
| 1 | A\|A |  |  | AA\|AA | distinguishable |
| 2 | A\|H |  | AB\|AA, AA\|AB |  | distinguishable |
| 3 | A\|B | AB\|AB |  |  | distinguishable |
| 4 | H\|A |  | AA\|BA, BA\|AA |  | distinguishable |
| 5 | H\|H | AB\|BA, BA\|AB |  | AA\|BB, BB\|AA | cannot distinguish DR and NR |
| 6 | H\|B |  | BB\|AB, AB\|BB |  | distinguishable |
| 7 | B\|A | BA\|BA |  |  | distinguishable |
| 8 | B\|H |  | BB\|BA, BA\|BB |  | distinguishable |
| 9 | B\|B |  |  | BB\|BB | distinguishable |

DR: double recombinant; SR: single recombinant; NR: non-recombinant

Now let the recombination frequency between the two loci be *r*. Thus, the probabilities of the 16 possible genotypes are given in Table A4, they are derived from the gamete probabilities (Table A5).

If we could distinguish all the classes (for instance, if the male and female parents had distinguishable alleles), then the estimator for *r* would be:

**Table 4:** Genotypes probabilities or expected frequencies of the 16 true genotypes.

| # | True<br>genotype | Type | $p(NR)$ | $p(DR)$ | $p(SR)$ | Sum | Name<br>class |
| --- | --- | --- | --- | --- | --- | --- | --- |
| 1 | AA\|AA | NR\|NR = NR | $(1 - r)^2/4$ | | | $(1 - r)^2/4$ | A |
| 2 | AA\|BB | NR\|NR = NR | $(1 - r)^2/4$ | | | $(1 - r)^2/4$ | B |
| 3 | BB\|AA | NR\|NR = NR | $(1 - r)^2/4$ | | | $(1 - r)^2/4$ | C |
| 4 | BB\|BB | NR\|NR = NR | $(1 - r)^2/4$ | | | $(1 - r)^2/4$ | D |
| 5 | AB\|AB | R\|R = DR | | $r^2/4$ | | $r^2/4$ | E |
| 6 | AB\|BA | R\|R = DR | | $r^2/4$ | | $r^2/4$ | F |
| 7 | BA\|AB | R\|R = DR | | $r^2/4$ | | $r^2/4$ | G |
| 8 | BA\|BA | R\|R = DR | | $r^2/4$ | | $r^2/4$ | H |
| 9 | AA\|AB | NR\|R = SR | | | $r(1 - r)/4$ | $r(1 - r)/4$ | I |
| 10 | AA\|BA | NR\|R = SR | | | $r(1 - r)/4$ | $r(1 - r)/4$ | J |
| 11 | BB\|AB | NR\|R = SR | | | $r(1 - r)/4$ | $r(1 - r)/4$ | K |
| 12 | BB\|BA | NR\|R = SR | | | $r(1 - r)/4$ | $r(1 - r)/4$ | L |
| 13 | AB\|AA | R\|NR = SR | | | $r(1 - r)/4$ | $r(1 - r)/4$ | M |
| 14 | AB\|BB | R\|NR = SR | | | $r(1 - r)/4$ | $r(1 - r)/4$ | N |
| 15 | BA\|AA | R\|NR = SR | | | $r(1 - r)/4$ | $r(1 - r)/4$ | O |
| 16 | BA\|BB | R\|NR = SR | | | $r(1 - r)/4$ | $r(1 - r)/4$ | P |
| Sum | | | $(1 - r)^2$ | $r^2$ | $2r(1 - r)$ | 1 | $n$ |

**Table A5:**
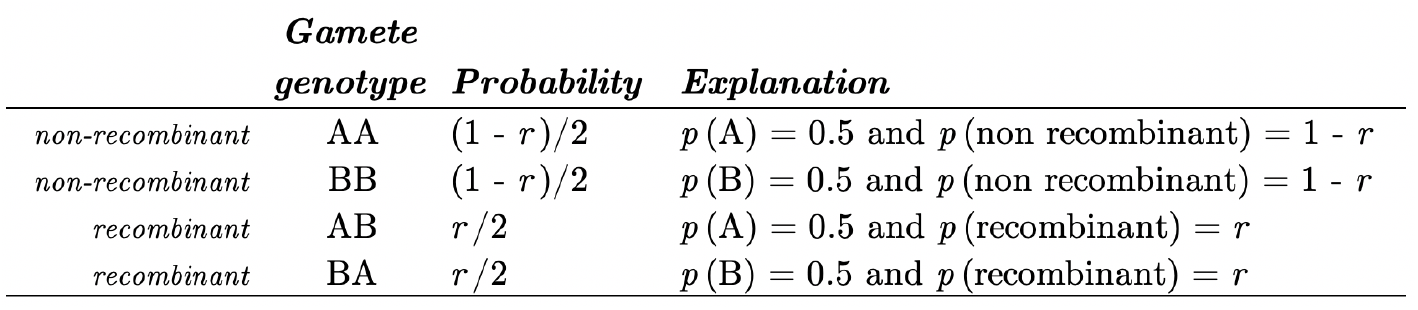
Gametes probabilities (male or female)

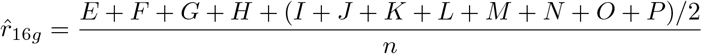

which is the estimator of the average of male and female recombination frequencies, with asymptotic variance

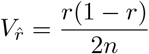

The expected frequency of SR-equivalents is 2r. This is because the F_**2**_ progeny results from two meioses, while in the backcross only one meiosis is observed. Note that we can estimate female and male recombination separately

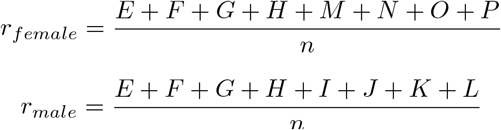

Now, consider the probability table for the 9 observable genotypes (Table A6).

We see that the male and female recombination frequencies cannot be estimated separately since the M, N, O and P classes are indistinguishable from the classes I, J, K and L. We can however estimate an average *r* in simply counting the observable recombinants:

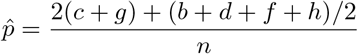

**Table A6:**
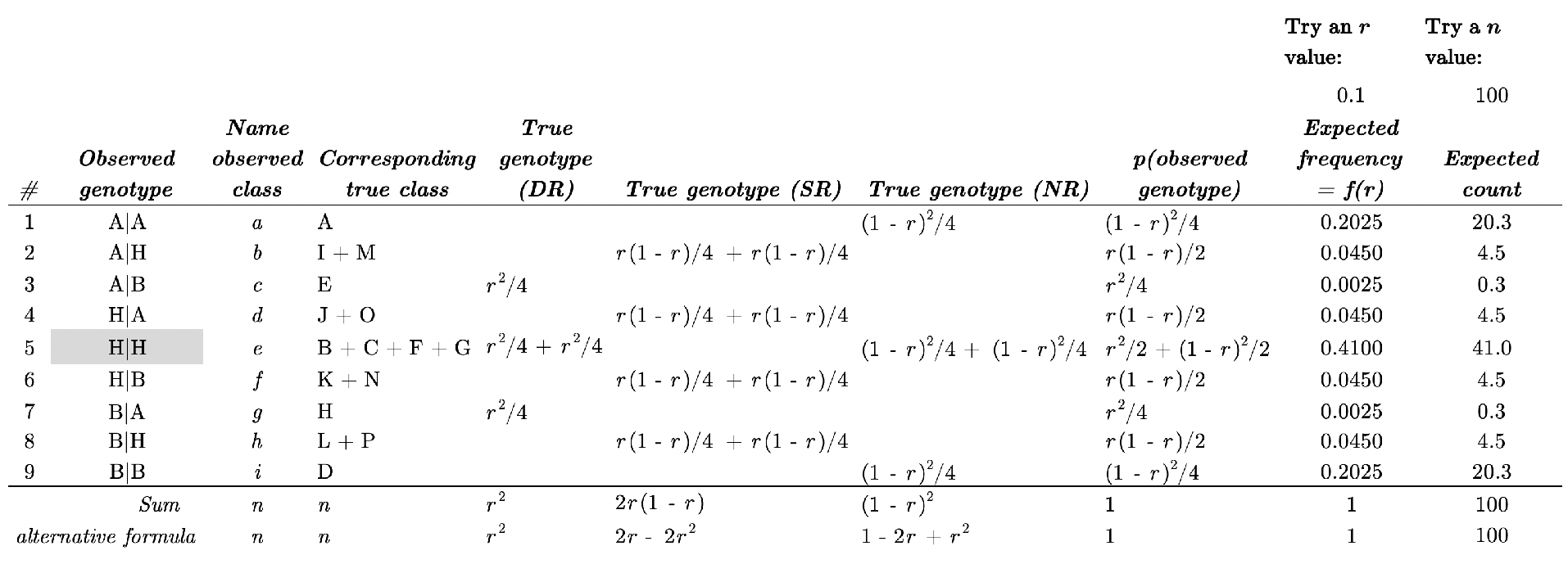
Genotypes probabilities or expected frequencies of the 9 observable genotypes.

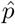 is the Benito & Gallego’s (2014) estimator, which is unbiased as shown in Table A7. Its variance is

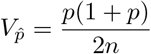

**Table A7:**
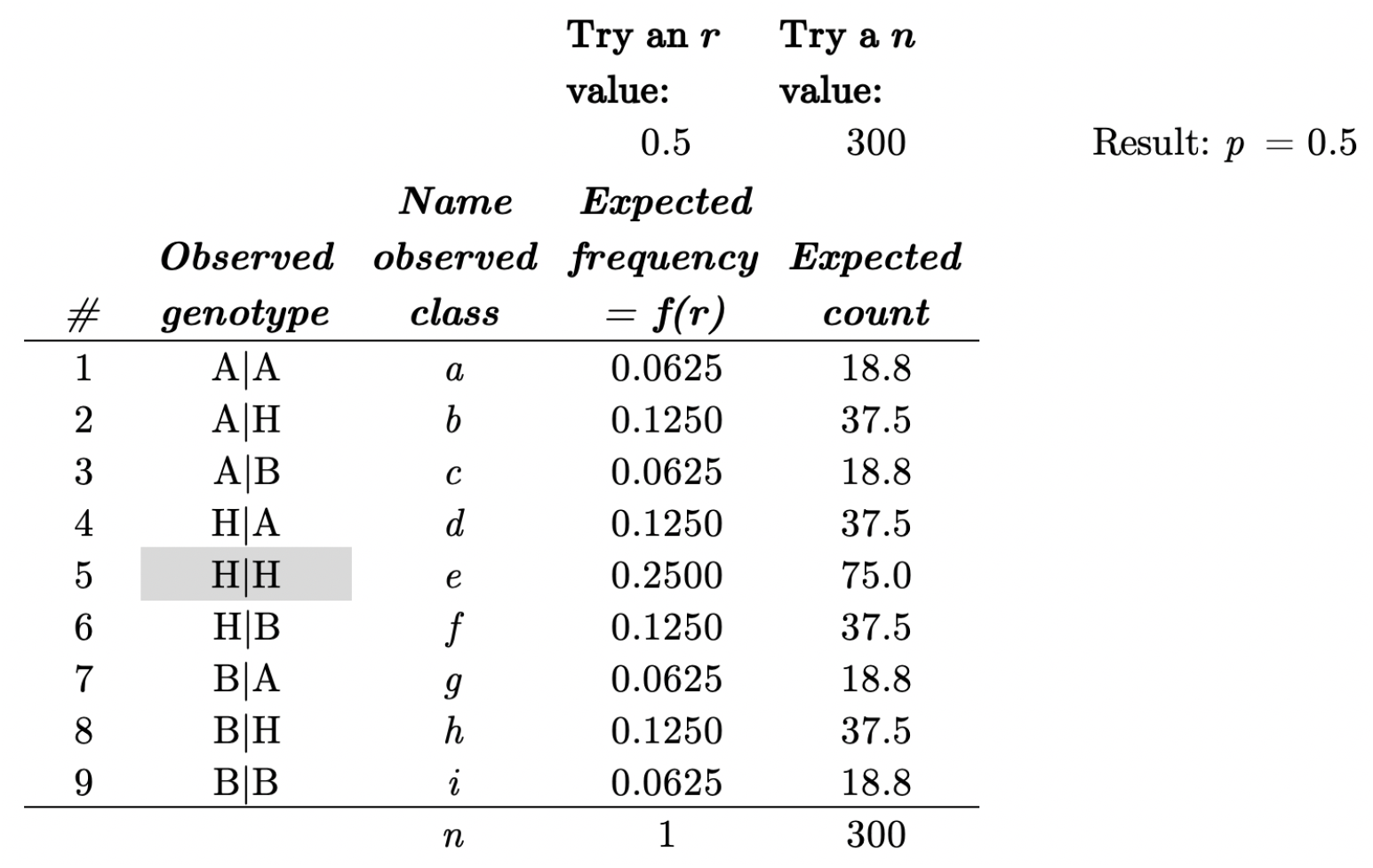
estimating r using the distinguishable classes only.

However, if ignoring the *H*|*H* class does not affect the informative sample size when *r* is small (< 0.3), this results in the loss of 12.5% of informative recombinants when *r* = 0.5.

This affects the variance of the estimator for *r* (Table A8).

The way to take the *H* | *H* class into account is to solve the equation

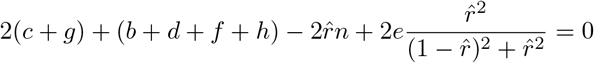

This can be attained using an iterative process like the Newton-Raphson or the EM algorithms. For instance, Mangin (1991) proposed the following EM:

**Table A8:**
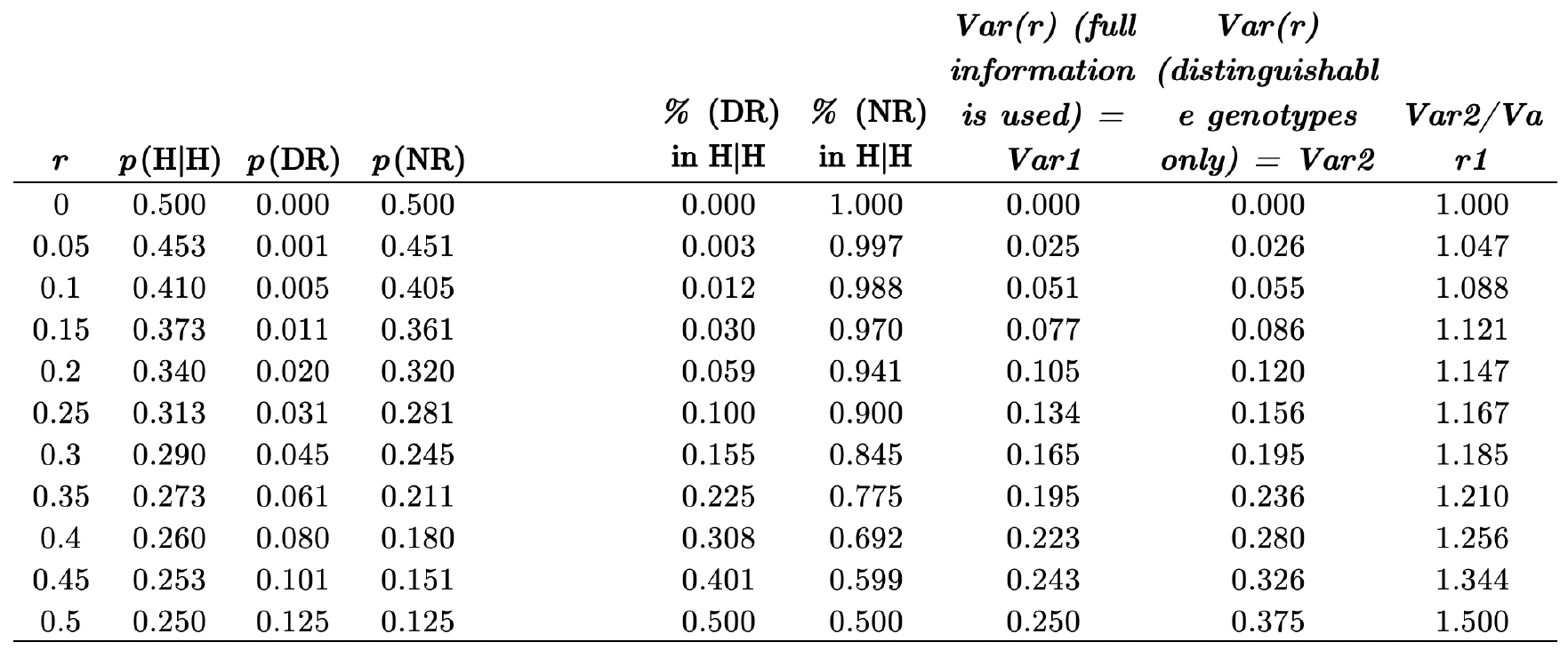
How do p(DR) and p(NR) within the H/H class vary with r?

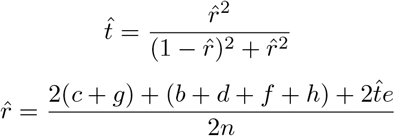

This provides, after convergence, a maximum-likelihood, unbiased and efficient estimator for *r*, with variance

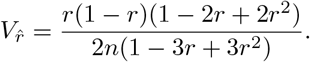

